# Study on the interference of *DicdsVg4* and *DicdsVgR* in egg formation in the ovary of *Diaphorina citri* (Kuwayama)

**DOI:** 10.1101/2022.04.15.488471

**Authors:** Hailin Li, Xiaoyun Wang, Biqiong Pan, Xialin Zheng, Wen Lu

## Abstract

The ovary is the key organ in insect female oviposition. For the continuation of populations in nature, the whole life cycle of female adults is centred on reproduction. *Vg* and *VgR* genes are important in determining the synthesis and transport of vitellogenin. Therefore, *Vg* and *VgR* gene expression in insect females affects the formation of mature eggs in the ovary, which is very important. RNAi interference of *Vg* and *VgR* genes may become an important means to control *D. citri* populations. A study found that the expression of the *DicVg4* and *DicVgR* genes was significantly affected by *DicdsVg4* and *DicdsVgR. DicVg4* and *DicVgR* gene expression interference could prevent the accumulation of vitellogenin in the ovaries of females, rendering ovaries unable to form mature eggs normally and leading to the production of abnormal eggs and nymphs. After knockdown of *DicVg4* and *DicVgR* gene expression, the proportion of ovarian mature eggs was only 8.33 ± 1.67% and 7.75 ± 0.5% in the 10-d developmental stage, the proportion of negative control group ovarian mature eggs was 13.19 ± 0.14% in the 10-d developmental stage, and the proportion of ovarian mature eggs in the interference group was 0.00 ± 0.00% in the 5, 15, 20, 25, and 30-d developmental stages. These results provide a theoretical basis for the use of mixed polygene interference technology to control damage from *D. citri* populations.

## INTRODUCTION

The insect ovary is the key organ in insect population mating and oviposition, and the numbers of female individuals and ovarian eggs determine whether the insect population can persist in nature for a long time (Bueno et al., 2020; Koji et al., 2020; Amano and Nomura, 2021; Gotoh and Sasaki, 2021). A study found that the morphological characteristics of the internal reproductive organs of females, especially ovaries, change dramatically in a short time from emergence to death for most insects (Chiang, 2020; Haddad et al., 2021; Ji et al., 2021). Because individual pest population ovary metabolism is more active, females can lay a large number of eggs, and the life span of some pest populations can be up to 2-6 months. A study found that ovarian characteristic changes in female pests are particularly remarkable during development (Chen et al., 2022). Therefore, systematic study of the development of pest ovaries is helpful to find appropriate methods to control pest populations.

Controlling ovary development can effectively control pest populations. A study found that interfering with the expression of female oviposition-related genes can effectively control female oviposition. For example, interfering with the expression of the *Vitellogenin* (*Vg*) gene and *Vitellogenin receptor* (*VgR*) gene can effectively hinder vitellogenin accumulation and mature egg formation in ovaries, which can control pest populations (Ibanez et al., 2017; Peng et al., 2020). At present, various pest ovarian structures have been studied and recorded in detail by anatomical observation and microcomputed tomography (Javier et al., 2019; Ignacio et al., 2019; Attardo et al., 2020; Men et al., 2020; Dantas et al., 2021; Wyber et al., 2021). RNA interference (RNAi) and CRISPR-associated protein 9 (CRISPR/cas9) based on molecular biology are more advanced genetic engineering technologies that provide novel ideas and means for controlling pest ovarian development (Abdullaha et al., 2019; Chen et al., 2019; Fandino et al., 2019; Hao et al., 2019; Liu et al., 2019; Lin et al., 2019; Lu et al., 2019a; Ortega and Killiny, 2019; Wang et al., 2019; Dias et al., 2020). For example, an investigation of the characteristics and functions of the *small heat shock protein* (*SHSP*) genes of *Tribolium castaneum* (Coleoptera: Tenebrionidae) showed that the *heat shock protein 18*.*3* (*hsp18*.*3*) gene was expressed in all stages of adult development and highly expressed in the early pupal stage and late adult stage, especially in the ovary and fat body of adults, and that reproductive capacity was significantly reduced by silencing the *Tchsp18*.*3* gene; knockout of the *Tudor* gene in *Bactrocera dorsalis* (Diptera: Trypetidae) led to a decrease in ovarian development, mating rate, and fecundity and interfered with the corresponding expression of downstream genes and the expression of primordial germ cells (Xie et al. 2019a; Xie et al., 2019b).

*Diaphorina citri* (Kuwayama) (Homoptera: Liviidae) is a citrus Huanglongbing (HLB) disease vector that often leads to disease outbreaks, and citrus orchards that are destroyed by HLB will not make profits in the next 10 years (Stockton et al., 2017). At present, a study found that *D. citri* adults can spread over long distances with high wind speeds in the autumn, and large-scale cultivation of Rutaceae crops all over the world has led to a continuous increase in individual populations and expansion of damage (Li et al., 2019; Li et al., 2020). To control HLB disease spread, RNA interference (RNAi) technology is applied to *Vg* and *VgR* gene expression in females to hinder the normal development of ovaries, which represents a new idea for using molecular biology technology to control *D. citri* population. A study found that egg production by females requires the mobilization of nutrients for egg cells in the ovary to produce vitellogenin (Lu et al. 2019b). Vg protein is synthesized in the fat body, secreted into the haemolymph, and then incorporated into developing oocytes through vitellogenin receptor (VgR)-mediated endocytosis (Tufail and Takeda, 2008). Therefore, RNAi technology was used to interfere with the expression of the female *Vg* and *VgR* genes and the formation of eggs as a feasible approach to control damage from *D. citri* populations. A study found that *Vg* gene expression can directly or indirectly affect the development of insect ovaries, and *VgR* gene expression can affect insect egg formation, oviposition behaviour, and other physiological activities of oviposition (Ibanez et al., 2017; Lu et al., 2019b). Differential gene expression analysis of *D. citri* females based on transcriptome sequencing screened out *Vitellogenin-1-like-1* (*Vg1*), *Vitellogenin-1-like-2* (*Vg2*), *Vitellogenin-2-like* (*Vg3*), *Vitellogenin-3-like* (*Vg4*), *Vitellogenin-like* (*Vg5*) and *VgR*, and this study provides a theoretical basis for the interference of *Vg* and *VgR* gene family members in *D. citri* females (Li et al., 2022). However, after interference with *D. citri* ovary *Vg* and *VgR* gene expression, the developmental characteristics of the ovary were not recorded in detail. Studies have only made a rough estimate of *D. citri* internal reproductive organ anatomy, identified basic ovarian morphology, and ovarian developmental processes have been systematically reported (Xiao et al., 2017; Guo et al., 2021). After interference with *D. citri Vg* and *VgR* gene expression, the developmental characteristics of the ovary should be thoroughly studied and recorded. It is worth noting that some studies have researched *D. citri* adult internal reproductive organs by micro computed tomography and anatomical means, but the accumulation process and mechanism of ovarian vitellogenin need further study (Ruan et al., 2012; Javier et al., 2021).

In this study, we tested whether *DicdsVg4* and *DicdsVgR* could effectively reduce the expression of the *DicVg4* and *DicVgR* genes and interfere with the formation of ovarian mature eggs. To answer our question, we examined the (1) stability of dsRNA in *M. odorifera* shoots; (2) *DicVg4* and *DicVgR* gene expression and ovarian development; and (3) changes in egg morphology and amount and numbers of malformed eggs and nymphs. The results can provide a theoretical basis for polygen mixed interference technology to control *D. citri* spread and harm in Rutaceae production areas worldwide.

## MATERIALS AND METHODS

### Insect rearing and collection

Adults of *D. citri* were collected from *Murraya odorifera* plants in the citrus orchard of Guangxi University (108.290°N, 22.849°E) and raised in five indoor cages (90 × 90 × 100 cm^3^). The 80-100 cm-tall *M. odorifera* plants were transferred to the insect cages as food for adults. Six hundred adults that emerged after 8-11 days were paired in the indoor cages as mentioned above, and the young shoots of *M. odorifera* seedlings were used as oviposition substrates. Adults of the offspring were obtained as experimental materials.

Males and females that emerged on the same day were collected using a 50 mL finger tube for extraction of total RNA, synthesis of dsRNA, and anatomical observation of internal reproductive organs. All experiments were conducted indoors at a temperature of 25 ± 1 °C, relative humidity of 75 ± 5%, and light:dark = 14:10 h.

### Delivery and stability of dsRNA

Before dissection, females were placed in liquid nitrogen to abolish their ability to move. Precooled 1× PBS (37 mM NaCl, 2.68 mM KCl, 8.1 mM Na_2_HPO_4_, 1.47 mM KH_2_PO_4_, pH 7.4) was added to alcohol-treated wax tablets, and the abdomens of *D. citri* females were immediately dissected under a stereoscope (SMZ800N, Nikon, JP). Micron insect anatomical needles were used to cut the side of the female abdomen and remove the exoskeleton on the back of the abdomen (YuRenTang, CN). The internal reproductive organs of the females were removed from the end of the abdomen. Micron insect needles were used to place the ovary in a 1.5 mL EP tube with PBS. EP tubes with ovaries were immediately frozen in liquid nitrogen and stored at -80 □ until RNA extraction. *D. citri* ovary total RNA was isolated using *TransZol* Up according to the manufacturer’s instructions (TransGen Biotech, CN). After the collected groups of six *D. citri* ovaries were ground using a pestle, *TransZol* was added to the EP tubes, and total RNA was extracted following the manufacturer’s instructions. After total RNA of *D. citri* ovaries was obtained, cDNA synthesis was carried out using the Primescript™ RT reagent kit with gDNA eraser (perfect real time) (TaKaRa, JP) following the manufacturer’s instructions.

*DicVg4* (477 bp) cDNA fragments were generated from the RNA of *D. citri* ovaries with the primer pair DicVg4 F3/R3 using 2×FastPfu Premix (TOLOBIO, USA). The purified PCR fragments were cloned into the pMD18T vector (DH5α, Thermo Fisher Scientific, USA). The resulting *DicVg4* plasmids were used as templates to generate *DicdsVg4* with the RNAsyn dsRNA kit (RNAsyn, CN). Synthesized dsRNA was quantitated by a Nanophotometer N80 Touch (Implen, GER) at 260 nm, and the integrity was analysed by agarose gel electrophoresis. The methods of *DicdsVgR* (346 bp) and *dsGPF* (726 bp) synthesis were the same as those for *DicdsVg4* (Table 1).

**Table 1.**
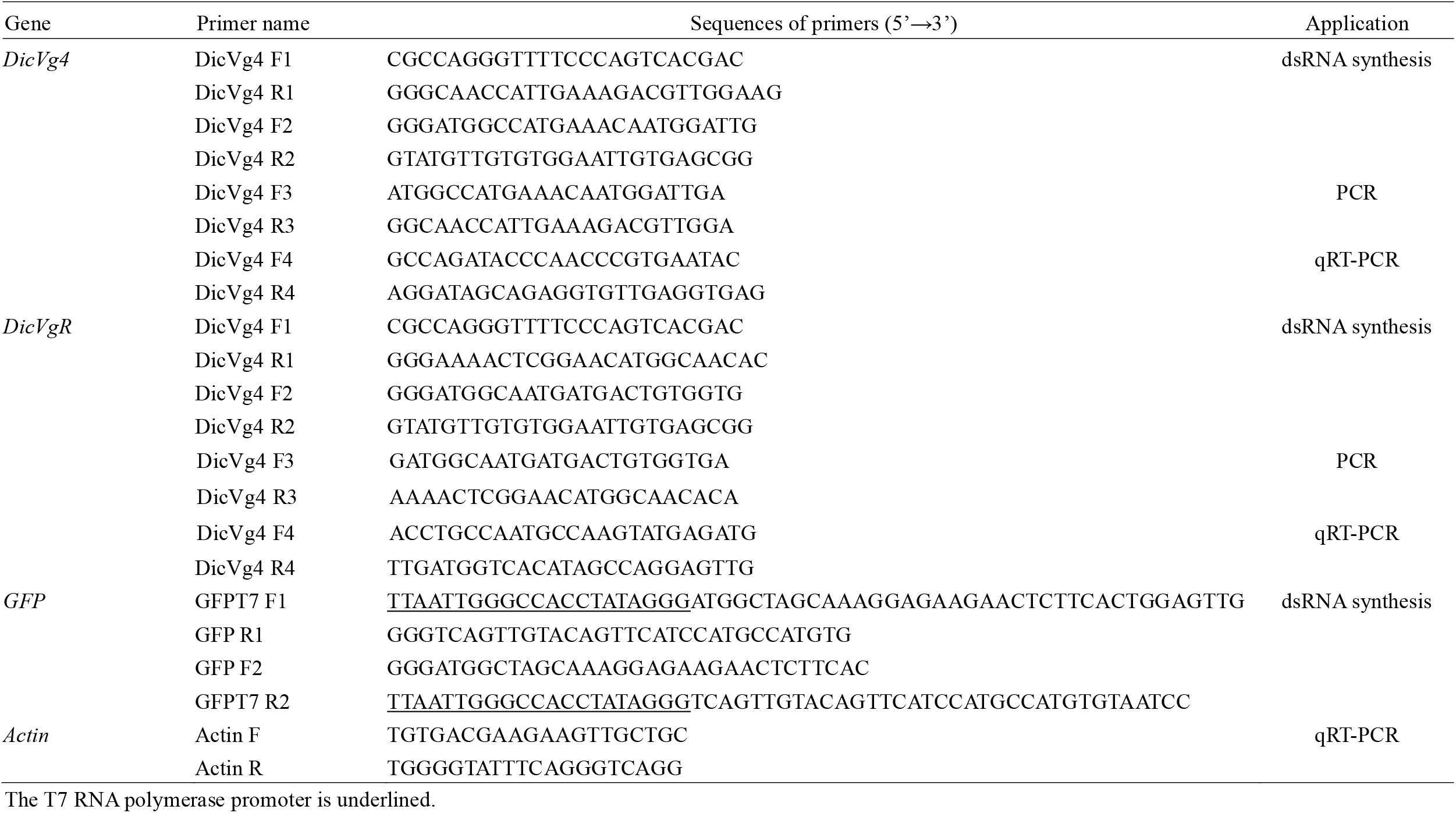
Oligonucleotide primer pairs used in this study.

Six *M. odorifera* shoots were immersed in a plastic vehicle containing *DicdsVg4* (1 ml) solution at a concentration of 10 ng/µl (Fig. 1). Individual *M. odorifera* shoot and leaf samples were collected daily from plastic vehicles, immediately frozen in liquid nitrogen and stored at -80 □ until RNA extraction. *M. odorifera* shoot and leaf total RNA was isolated using *TransZol* Up according to the manufacturer’s instructions. cDNA was synthesized from *M. odorifera* leaf total RNA, and the *DicVg4* gene was cloned into a cDNA template. The stability of *DicdsVg4* at 6 d was detected by gel electrophoresis. The stability detection method of *DicdsVgR* was the same as that for *DicdsVg4*.

**Fig. 1.**
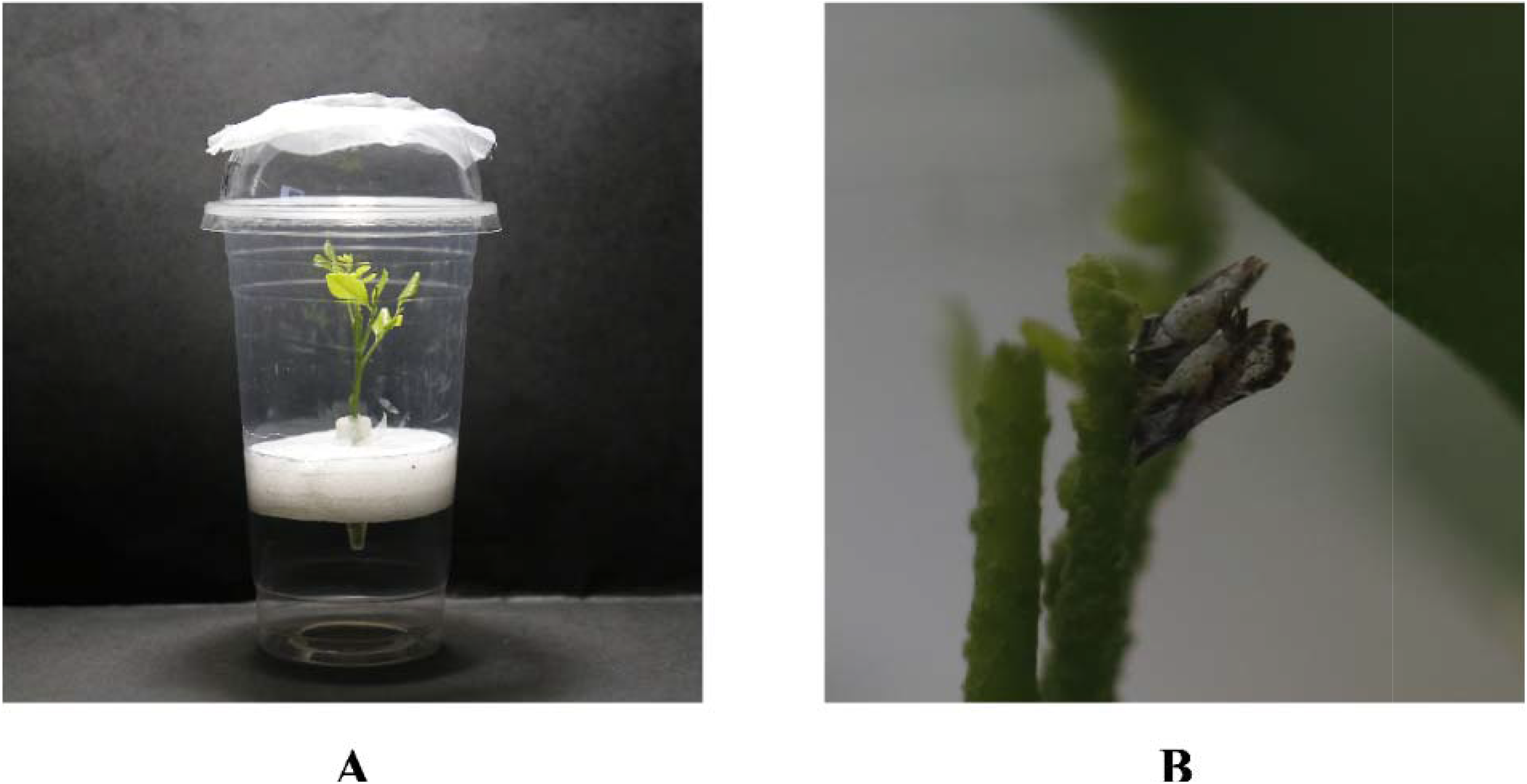
Insect rearing device. (A) Plastic vehicle with *M. odorifera*. (B) Mating of male and female adults.

### Anatomy of the *D. citri* ovary and quantitative real-time PCR (qRT–PCR) analysis

The collected *D. citri* males and females were placed in a plastic vehicle with *M. odorifera* shoots, 1 mL of 10 ng/µl sterile water containing *DicdsVg4* was placed into the plastic vehicle with an EP tube, and 1-2 *M. odorifera* shoots were added (Fig. 1). Fifteen pairs of males and females were raised in each plastic vehicle, and a total of 6 plastic vehicles were prepared. Females in one plastic vehicle were dissected every 5 d with micron insect needles (YuRenTang, CN). The female internal reproductive organs are shown in Fig. 2. Micron insect needles were used to place the female’s internal reproductive organs on a glass slide with PBS, and a stereoscope was used to observe and record the ovarian anatomy. During the experiment, the morphological characteristics of eggs oviposited by females and nymphs hatched from eggs were observed and recorded by stereoscopy, and malformed eggs and nymphs required special observation and recording. The anatomical method of feeding *DicdsVgR* and *dsGFP* females was the same as that for *DicdsVg4*. All treatments were repeated 3 times (Table 1).

**Fig. 2.**
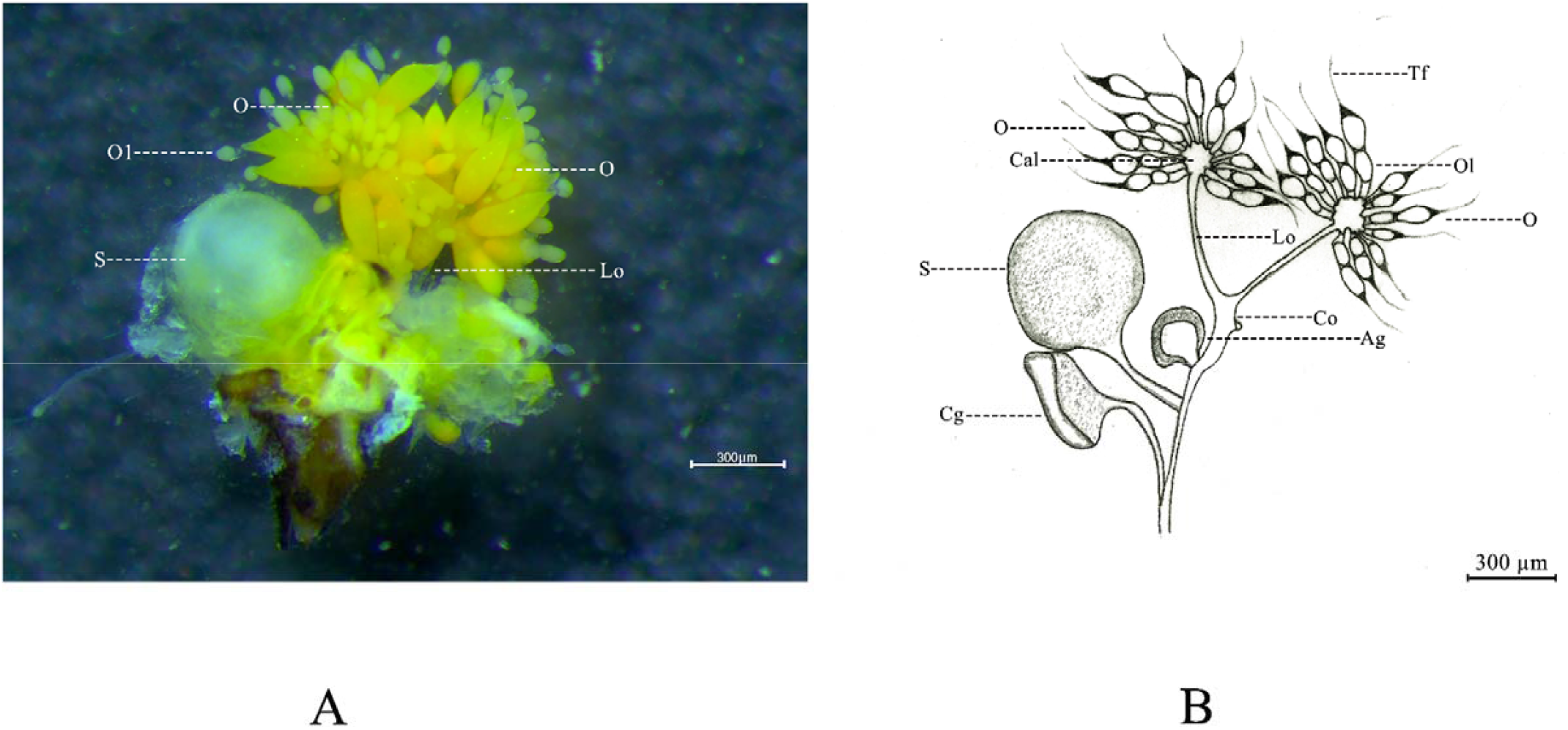
Ovarian morphology of *D. citri* females. (A) Ovarian morphology. (B) Hand drawing of ovarian morphology. Ag, accessory glands; Cal, calyx; Cg, colleterial gland; Co, common oviduct; Lo, lateral oviduct; O, ovary; Ol, ovariole; S, spermatheca; Tf, terminal filament.

The size of the egg was measured by using stereoscope measurement software. The length and width of eggs were measured, the number of ovary eggs was counted, and a microscopic measurement system matched with the stereoscope was used to record and observe the changes in the ovary (SMZ800N, Nikon, JP). Photoshop (Adobe Systems, US) was used to process the pictures (Fig. 3). After obtaining adult egg length data, ovary egg number data, and relative gene expression data, SPSS 17.0 (IBM, US) was used to analyse the egg length data and ovary egg number data for 1-30 d.

**Fig. 3.**
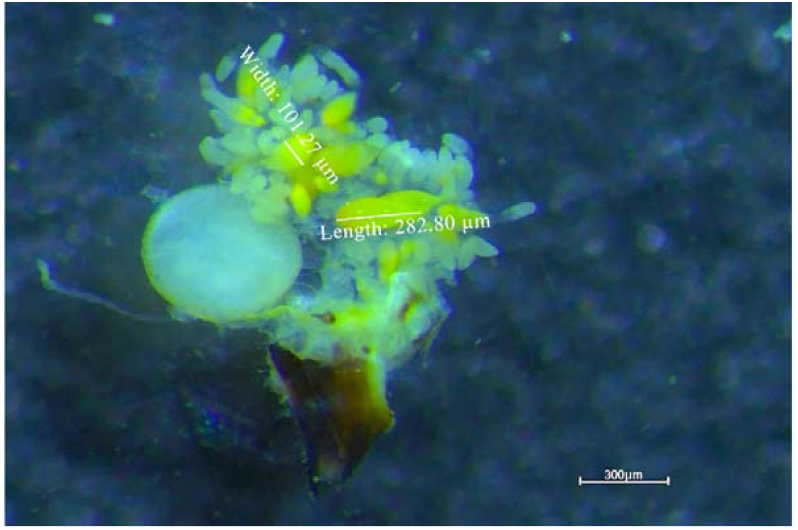
Measurement of *D. citri* eggs. The length and width of the eggs were measured.

After the expression of the *DicVg4* gene in the ovary was interfered with by *DicdsVg4*, anatomic ovaries were used for *DicVg4* gene expression analysis, including those of 5-, 10-, 15-, 20-, 25-, and 30-day-old developing females. The anatomic ovaries were stored at -80 □ until RNA extraction. Anatomic ovary total RNA was isolated using *TransZol* Up according to the manufacturer’s instructions. CDNA was synthesized from the anatomic ovary total RNA. The primer pairs used for qRT–PCR are shown in Table 1. qRT-PCRs were carried out using 2×Q3 SYBR qPCR Master Mix (High Rox) (TOLOBIO, USA). All reactions were performed with the QuantStudio™ Real-Time PCR system (Applied Biosystems, USA). The expression analysis of *DicdsVgR* and *dsGFP* was the same as that of *DicdsVg4*. All treatments were repeated 3 times. Relative gene expression data from real-time quantitative PCR were analysed with the 2^-^ΔΔCT method.

## RESULTS

### Stability of dsRNA in *M. odorifera* shoots

Gel electrophoresis was used to examine the stability of *DicdsVg4* and *DicdsVgR* in *M. odorifera* shoots at 6 d of treatment, revealing clear bands at the 1-2 d stage, which showed that *DicdsVg4* and *DicdsVgR* still maintained a relatively complete double-stranded structure at 1-2 d. After *D. citri* females fed on the *M. odorifera* shoots for 1-2 d, *DicdsVg4* and *DicdsVgR* smoothly entered the female body. After 3-6 d, the gel electrophoresis bands became blurred, which indicates that *DicdsVg4* and *DicdsVgR* began to decompose at this stage, but *DicdsVg4* gel electrophoresis revealed traces of bands that showed that the structure of some *DicdsVg4* and *DicdsVgR* remained complete at the 3-6 d stage. These results confirm that *DicdsVg4* and *DicdsVgR* could be absorbed by the *M. odorifera* shoots and be ingested by the *D. citri* females during the feeding process (Fig. 4).

**Fig. 4.**
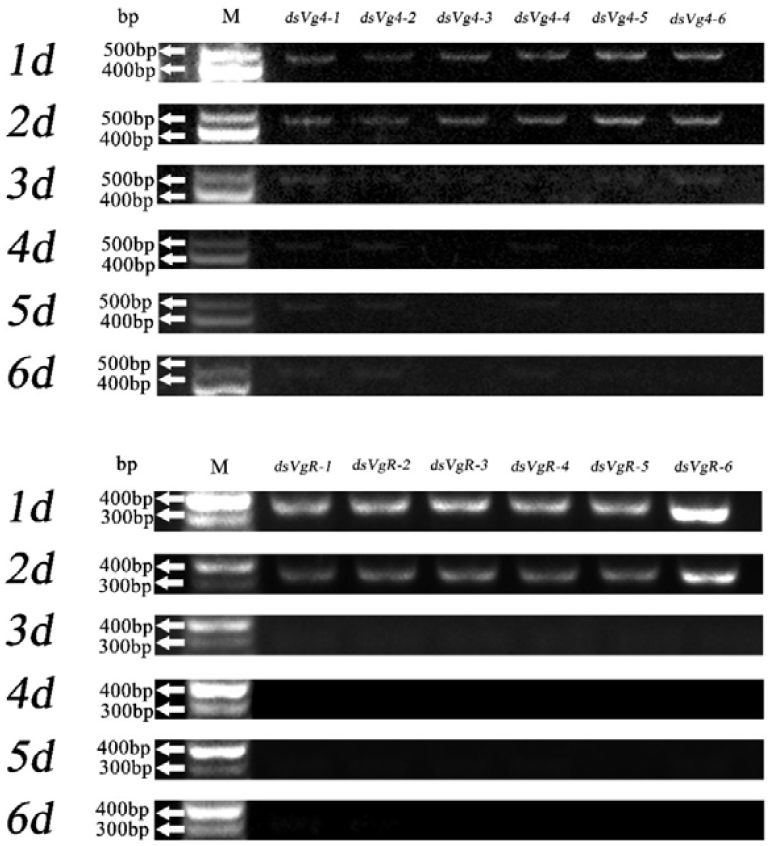
After *DicdsVg4* and *DicdsVgR* were absorbed by tender *M. odorifera* shoots, gel electrophoresis was used to detect the presence time of *DicdsVg4* and *DicdsVgR*.

### *DicVg4* and *DicVgR* gene expression and ovarian development

After *D. citri* females were fed *DicdsVg4*, the expression of the *DicVg4* gene decreased from 1-30 d. The expression of the *DicVg4* gene was decreased most significantly at 15 d, and the relative expression decreased 99.06% (P<0.05; Tukey test). However, the expression of the *DicVg4* gene was increased abnormally at 20 d, and the relative expression increased by 143.65% (P<0.05; Tukey’s test) (Fig. 5). The results showed that at the 20-d developmental stage, females had strong immune regulation of *DicdsVg4*. By dissecting the internal reproductive organs of females with interference of *DicVg4* gene expression, we found that the number of eggs in the ovary of the *DicdsVg4* interference group was lower than that in the ovary of the negative control group at 1-30 d. Ovaries of the negative control group females began to form a large number of eggs from 15 d, and a certain number of eggs could be observed in the ovaries at 15-30 d. Fewer mature and immature eggs were observed in the ovaries of the interference group, and a large number of eggs had not formed in the ovaries. The results showed that *DicdsVg4* interference with *DicVg4* gene expression and ovarian development had a certain effect.

**Fig. 5.**
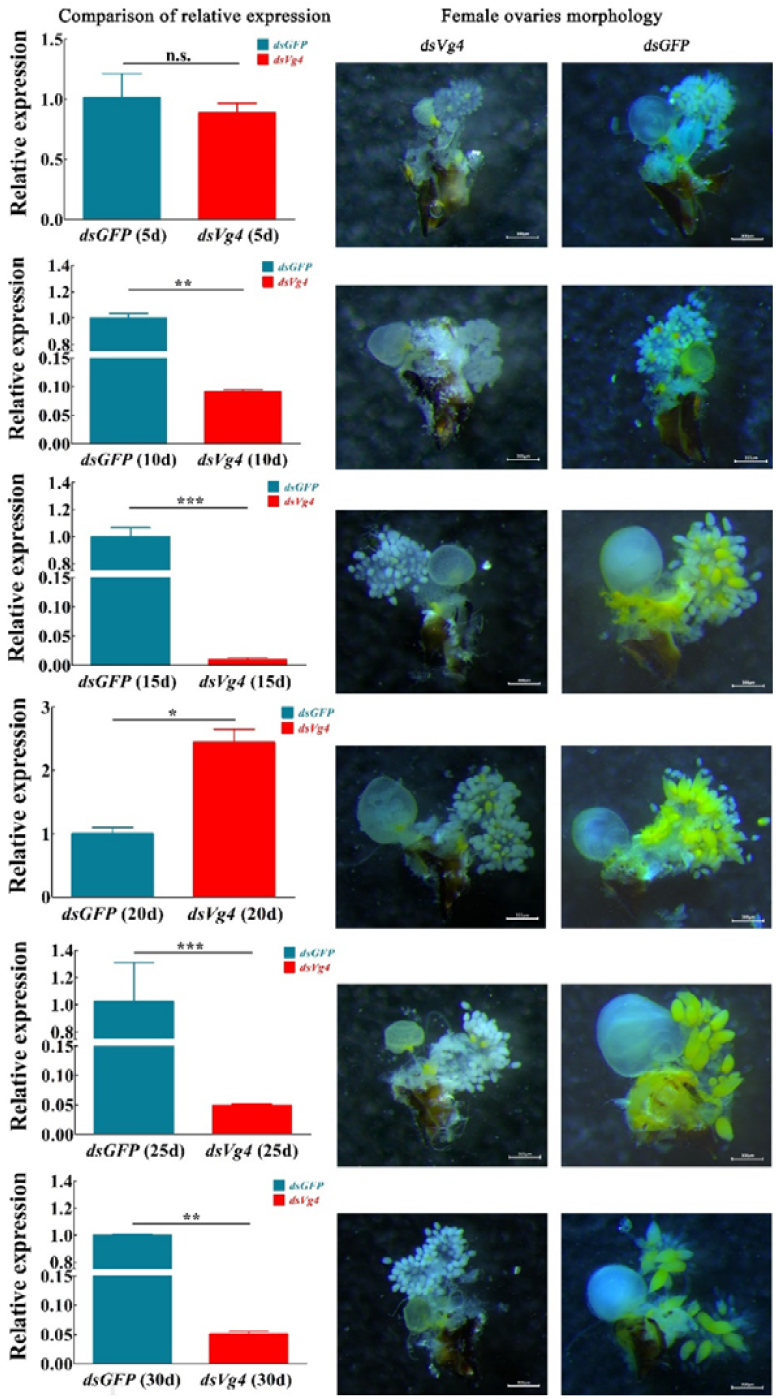
After *DicVg4* gene expression in *D. citri* females was knocked down, the relative expression of the *DicVg4* gene from 1-30 d and the developmental characteristics of the ovary were determined. The *DicdsVg4* gene interference group was compared with the negative control group. The length of the scale bar is 300 µm.

After *D. citri* females were fed *DicdsVgR*, the expression of the *DicVgR* gene decreased from 1-30 d. The expression of the *DicVgR* gene decreased most significantly at 25 d, and the relative expression decreased 99.57% (P<0.05; Tukey test). The expression of the *DicVgR* gene increased abnormally at 20 d, and the relative expression increased 256.93% (P<0.05; Tukey’s test) (Fig. 6). Females showed strong immune regulation of *DicdsVgR* at the 20-d developmental stage. By dissecting the internal reproductive organs of females with interference of *DicVgR* gene expression, it was found that the number of eggs in the ovary of the interference group was lower than that of the negative control group at 1-30 d, similar to the findings for the *DicdsVg4* interference group. Fewer mature and immature eggs were observed in the ovaries of the interference group, a large number of eggs had not formed in the ovaries, and fewer ovarian eggs were more obvious than in the *DicdsVg4* interference group. The results showed that the VgR receptor was important for the accumulation of Vg protein in the ovary.

**Fig. 6.**
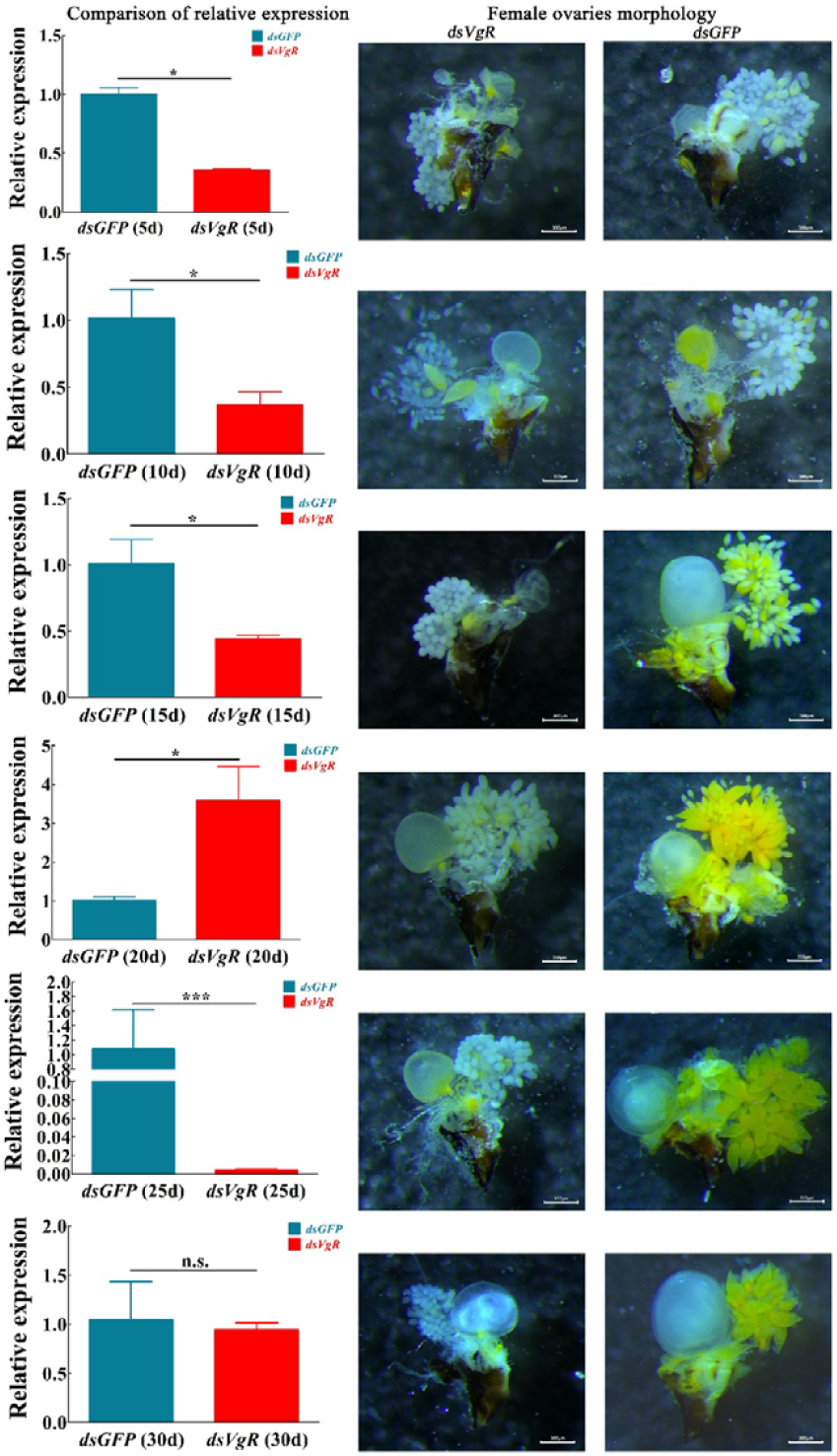
After *DicVgR* gene expression in *D. citri* females was knocked down, the relative expression of the *DicVgR* gene from 1-30 d and the ovarian development characteristics of females were determined. The *DicdsVgR* interference group was compared with the negative control group. The length of the scale bar is 300 µm.

### Changes to egg morphology and numbers of eggs, malformed eggs and nymphs

*DicVg4* and *DicVgR* gene expression in *D. citri* females was interfered with, resulting in a relatively small number of mature ovarian eggs after 1-30 d of development (Fig. 5, Fig. 6, Table 2). At the 5-d developmental stage, the ovaries of the interference group and negative control group did not form eggs. Only at the 10-d developmental stage, *DicdsVg4* and *DicdsVgR* interference group mature eggs were found in the dissected ovaries. At the 10-d developmental stage, *DicdsVg4* interference group eggs showed an average maximum length of 136.30 ± 15.55 µm, an average maximum egg width of 62.58 ± 6.30 µm, maximum number of mature eggs of 1.03 ± 0.37, and maximum percentage of mature eggs of 8.33 ± 1.67%. At the 10-d developmental stage, the *DicdsVgR* interference group showed an average maximum egg length of 123.32 ± 30.03 µm, average maximum egg width of 61.62 ± 6.59 µm, maximum number of mature eggs of 0.75 ± 0.25 grain, and maximum mature egg percentage of 7.75 ± 0.50%. Th egg size and mature egg proportion of *DicdsVg4* and *DicdsVgR* interference groups were lower than those of the negative control group at the 10-d developmental stage. At the 15-30 d developmental stage, mature eggs of the *DicdsVg4* and *DicdsVgR* interference groups were not observed, and only a few immature eggs were observed in the ovarioles. The negative control group average maximum egg length reached 293.71 ± 9.74 µm, and the average maximum egg width reached 124.24 ± 18.77 µm at the 25-d developmental stage, while the number of mature eggs reached a maximum of 13.25 ± 0.75 and the mature egg percentage reached a maximum of 52.91 ± 19.29% at the 20-d developmental stage. The egg size and mature egg proportion of the negative control group were higher than those of the *DicdsVg4* and *DicdsVgR* interference groups at the 15-30 d developmental stage. The results showed that after *DicVg4* and *DicVgR* gene expression in *D. citri* females was knocked down, the ability for vitellogenin accumulation in the ovary was decreased significantly, and insects were unable to form a certain number of mature eggs normally.

**Table 2.** The length and width, number, proportion of mature eggs in ovary of *D. citri* from 1-30 d. Data of the same day were compared vertically

| Day (d) | Different treatment | Egg |  |  |  |
| --- | --- | --- | --- | --- | --- |
|  |  | Length (μm) | Width (μm) | Number of mature eggs (grain) | Mature eggs percentage (%) |
| 5d | *DicdsVg4* | 0.00±0.00a | 0.00±0.00a | 0.00±0.00a | 0.00±0.00a |
|  | *DicdsVgR* | 0.00±0.00a | 0.00±0.00a | 0.00±0.00a | 0.00±0.00a |
|  | *dsGFP* | 0.00±0.00a | 0.00±0.00a | 0.00±0.00a | 0.00±0.00a |
| 10d | *DicdsVg4* | 136.30±15.55a | 62.58±6.30a | 1.03±0.37a | 8.33±1.67a |
|  | *DicdsVgR* | 123.32±30.03a | 61.62±6.59b | 0.75±0.25a | 7.75±0.5a |
|  | *dsGFP* | 142.67±45.99b | 66.51±21.81c | 1.29±0.46b | 13.19±0.14b |
| 15d | *DicdsVg4* | 47.99±3.99b | 28.02±3.54b | 0.00±0.00b | 0.00±0.00b |
|  | *DicdsVgR* | 0.00±0.00c | 0.00±0.00c | 0.00±0.00b | 0.00±0.00b |
|  | *dsGFP* | 249.16±50.81a | 108.29±11.35a | 5.33±1.33a | 16.08±5.80a |
| 20d | *DicdsVg4* | 71.26±13.52b | 41.63±4.97b | 0.00±0.00b | 0.00±0.00b |
|  | *DicdsVgR* | 79.78±9.61b | 46.06±8.52b | 0.00±0.00b | 0.00±0.00b |
|  | *dsGFP* | 285.83±37.97a | 109.52±8.12a | 13.25±0.75a | 52.91±19.29a |
| 25d | *DicdsVg4* | 75.17±9.90b | 43.79±6.99b | 0.00±0.00b | 0.00±0.00b |
|  | *DicdsVgR* | 0.00±0.00c | 0.00±0.00c | 0.00±0.00b | 0.00±0.00b |
|  | *dsGFP* | 293.71±9.74a | 124.24±18.77a | 11.38±0.88a | 23.36±4.47a |
| 30d | *DicdsVg4* | 0.00±0.00c | 0.00±0.00c | 0.00±0.00b | 0.00±0.00b |
|  | *DicdsVgR* | 70.64±8.87b | 43.96±0.24b | 0.00±0.00b | 0.00±0.00b |
|  | *dsGFP* | 286.91±37.67a | 115.77±7.93a | 9.33±3.00a | 18.91±6.42a |

Means followed by different small letter in columns are significantly different, Tukey test was used for analysis of variance (ANOVA followed by SPSS, P<0.05).

After *DicVg4* and *DicVgR* gene expression of *D. citri* females was interfered with, abnormal eggs and nymphs were found on the *M. odorifera* shoots (Fig. 7). Females of the negative control group produced normal eggs, which were shaped like water droplets, and the egg surfaces were yellow and smooth. The abnormal egg surfaces of the *DicdsVg4* and *DicdsVgR* interference groups were shrivelled and brown compared with normal eggs. Normal nymphs of the negative control group had smooth backs and yellow bodies, abnormal nymphs of the *DicdsVg4* and *DicdsVgR* interference groups were wrinkled, the nymph bodies were brownish yellow, and some nymph bodies of the *DicdsVg4* interference group were translucent. All the abnormal eggs did not hatch, and abnormal nymphs showed abnormal characteristics at the 1-2 instar stage. The results showed that the abnormal expression of the *DicVg4* and *DicVgR* genes in females could lead to distortions of eggs and nymphs.

**Fig. 7.**
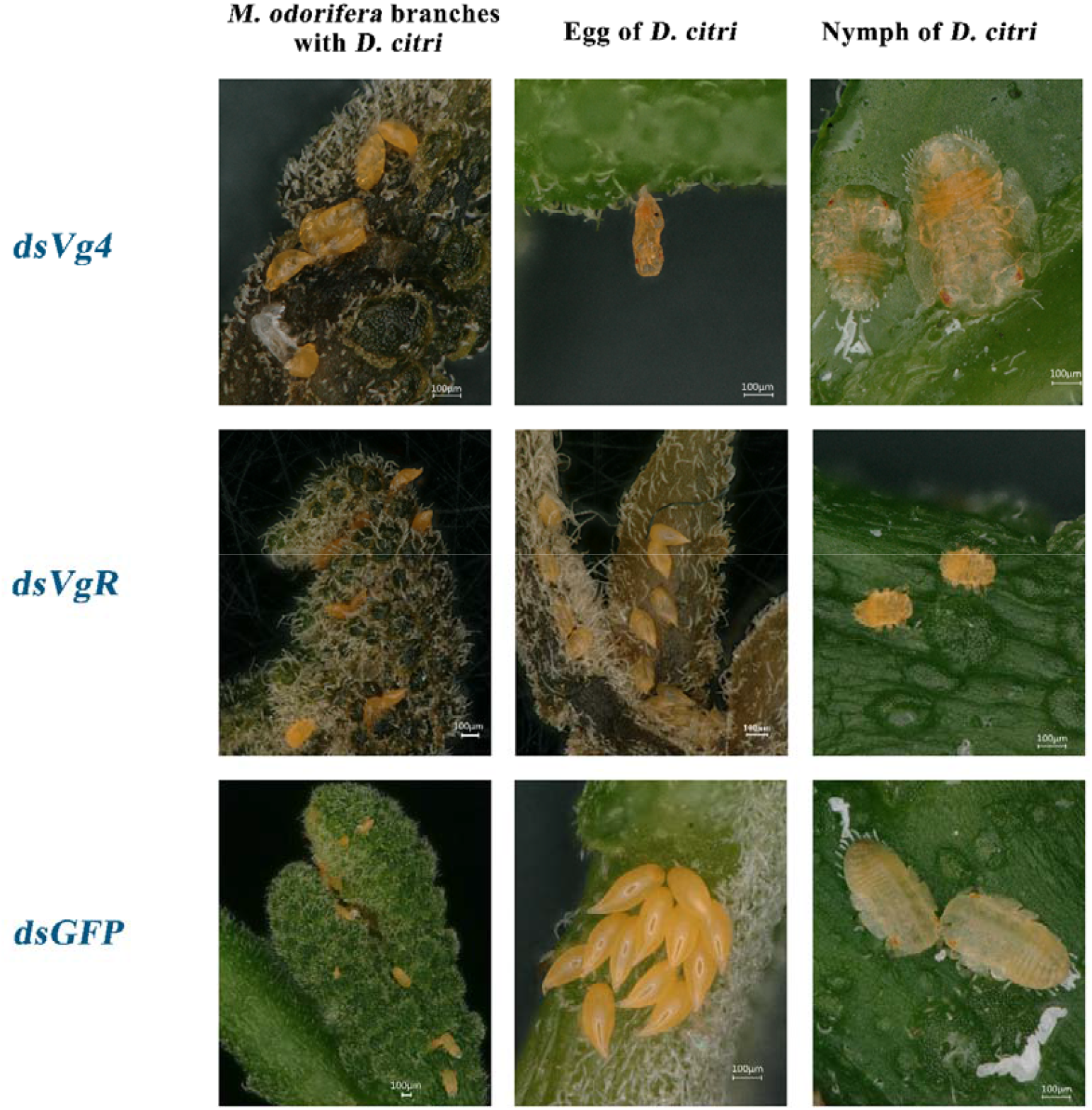
Malformed egg and nymph of *D. citri* females. *DicdsVg4* and *DicdsVgR* are the interference groups, and *dsGFP* is the negative control group.

## DISCUSSION

The results showed that *DicdsVg4* and *DicdsVgR* were very suitable for reducing numbers of *D. citri* ovarian eggs. The anatomy of the ovary showed that the expression of the *DicVg4* and *DicVgR* genes interfered with the decrease in Vg4 protein in the ovary, and the number of ovarian mature eggs also decreased accordingly. This shows that the Vg4 protein is very important for the formation of ovarian mature eggs. However, the functional differences between other members of the *D. citri Vg* gene family and the *Vg4* gene need to be further studied. This research suggested that *DicdsVg4* and *DicdsVgR* can effectively inhibit the expression of *DicVg4* and *DicVgR* genes, which can provide a theoretical basis for future research on the application of *DicdsVg4* and *DicdsVgR*.

Based on the stability of the dsRNA in *M. odorifera* shoots, it was found that delivery of dsRNA to a pest insect is easy through plant absorption of dsRNA. DsRNA can exist in liquid form under certain environmental conditions for a period of time. A study found that after leaf tissue absorbed *dsGFP*, the leaf tissue tested positive for the presence of *dsGFP* 24 h post-treatment and remained positive up to 40 days post-treatment (Andrade and Hunter, 2016). The gel electrophoresis of *DicdsVg4* and *DicdsVgR* also showed that *DicdsVg4* and *DicdsVgR* could exist for 2-5 days. Therefore, dsRNA is very suitable for controlling populations of *D. citri* and can be used to replace some chemical pesticides (Ghosh et al., 2018).

The results showed that *DicVg4* and *DicVgR* gene expression was significantly affected by *DicdsVg4* and *DicdsVgR*. At the 1-30 d developmental stage, the relative expression of the *DicVg4* and *DicVgR* genes was detected at 6 time points, showing that the relative expression of genes decreased overall, but the relative expression of the *DicVgR* gene abnormally increased 256.93% at the 20-d developmental stage. This phenomenon may have been caused by the immune regulation of cells by dsRNA. Previous studies indicated that dsRNA-degrading enzymes (dsRNases) have been recognized as important factors in reducing RNA interference efficiency in different insect species and lead to an increase in gene expression interference (Garbutt et al., 2013; Chen et al., 2021). Usually, the secretion of dsRNA-degrading enzymes is only part of the insect immune response (Swevers et al., 2013). However, the immune response induced by dsRNA and the related mechanisms need further research.

Our previous study showed that the *Vg* (*vitellogenin*) gene and *VgR* (*vitellogenin receptor*) gene family have a decisive impact on ovary development (Li et al., 2022). The results from the current study also showed that the number of mature eggs in the ovary decreased significantly after *DicVg4* and *DicVgR* genes interfered with *DicdsVg4* and *DicdsVgR*. This result suggests that the *DicVg4* gene is closely related to the formation of mature ovary eggs. The function of the *DicVgR* gene is different from that of the *DicdsVg4* gene, and the *DicVgR* gene indirectly affects the accumulation of the Vg protein and the formation of mature eggs (Yao et al., 2018). The deletion of the VgR receptor can lead to insufficient accumulation of Vg4 protein in the ovary, so after interference with *DicVg4* and *DicVgR* gene expression, the characteristics of *DicdsVg4* and *DicdsVgR* interference group ovarian development were similar. Although our previous study found that the *DicVg* gene family has six genes with different functions (Li et al., 2022), the functions of the other five *DicVg* genes except for *DicVg4* are still unknown. Furthermore, the proportion of different Vg proteins transported by the VgR receptor is also unclear and needs further exploration.

In this study, abnormal eggs and nymphs in the *DicdsVg4* and *DicdsVgR* interference groups were observed for the first time. This result suggested that interference of the *DicVg4* and *DicVgR* genes of *D. citri* females not only affect the formation of ovarian mature eggs but also affect the phenotypic characteristics of eggs and nymphs. This could be attributed to interference of *DicVg4* and *DicVgR* so that ovarian development was disturbed in addition to decreased fecundity and egg hatchability (Moriyama et al., 2016; Rika et al., 2019; Zhang et al., 2021; Chen et al., 2022). Therefore, our results suggest that *DicdsVg4* and *DicdsVgR* are very suitable for controlling populations of *D. citri*. Previous studies reported that the effect of dsRNA combined with special nanomaterials may be better than some chemical pesticides against *D. citri* populations (Zhang et al., 2010; Cong et al., 2015; Zhang et al., 2015; Li et al., 2016; Zhen et al., 2018; Kim et al., 2019; Zhang et al., 2021). However, how to use dsRNA to achieve the best effect in controlling *D. citri* needs further study.

## Conclusions

In summary, the expression of the *DicVg4* and *DicVgR* genes was significantly affected by *DicdsVg4* and *DicdsVgR. DicVg4* and *DicVgR* gene expression interference can prevent the accumulation of vitellogenin in the ovary, which causes ovaries to be unable to form mature eggs normally and leads to the production of abnormal eggs and nymphs.

## Acknowledgements

The authors acknowledge experimental support from Hui-Li Ou Yang, Ran-Ran Su, Bi-Qiong Pan, and the large-scale instrument sharing platform of Guangxi University. **Competing interests**

The authors declare no competing or financial interests.

## Author contributions

Conceptualization: H.L.Li.; Methodology: H.L.Li., X.Y.Wang.; Validation: H.L.Li.; Formal analysis: H.L.Li., X.Y.Wang., B.Q.Pan., X.L.Zheng., W.Lu.; Investigation: H.L.Li.; Writing-original draft: H.L.Li., X.Y.Wang.; Writing-review & editing: H.L.Li., X.Y.Wang., W.Lu.; Visualization: H.L.Li., X.Y.Wang., W.Lu.; Funding acquisition: W.Lu.

## Funding

This work was financially supported by a grant from the Modern Agricultural Industry

Technology System Guangxi Innovation Team (Citrus) (grant no. nycytxgxcxtd-05-03) in China.

---

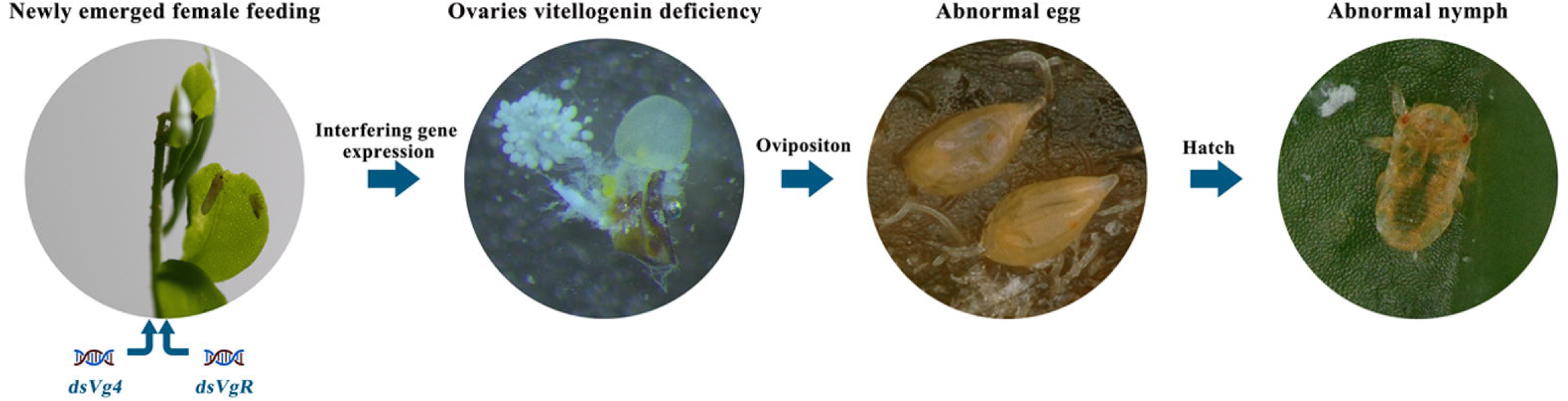

## Notes

### Competing Interest Statement

The authors have declared no competing interest.

